# DCS Tools: A high-performance, resource-efficient and scalable computing suite for population-scale genomic analysis and data compression

**DOI:** 10.64898/2026.03.13.711253

**Authors:** Chun Gong, Dong Yuan, Zijian Zhao, Xin Li, Yuxin Chen, Qi Yang, Ruiwen Wan, Shengkang Li, Yong Zhang

## Abstract

The advent of population-scale genomics has created significant bottlenecks in computational infrastructure and storage. Traditional BWA-GATK Best Practices require massive resources, while most existing acceleration solutions depend on specialized hardware like GPUs or FPGAs, increasing costs and limiting deployment. To address this, we developed DCS Tools, a high-performance suite optimized for standard CPU architectures. DCS Tools processes a 30X WGS sample from raw FASTQ to variant calls in just 1.79 hours on a 32-thread instance—a 16-fold speedup over traditional pipelines without additional hardware. The suite integrates DPGT for million-scale joint calling and introduces SeqArc and VarArc, which reduce storage footprints by up to 80% (FASTQ) and 66% (VCF) compared to GZIP. DCS Tools offers a cost-effective, hardware-agnostic solution for petabyte-scale genomic analysis.

## Statement of Need

The landscape of genomic research has shifted from individual sample analysis to massive cohort studies involving hundreds of thousands of individuals. Projects such as the UK Biobank, the China Kadoorie Biobank (CKB), and various agricultural “Pan-Genome” initiatives generate raw sequencing data at a scale that challenges the limits of modern data centers. In these contexts, the efficiency of the primary analysis pipeline—comprising Quality Control (QC), sequence alignment, and variant calling—directly dictates the feasibility and cost of the research.

Typically, large-scale cohort studies follow a three-stage framework: individual sample variant calling, multi-sample joint calling, and downstream population genetic analysis. This study focuses on the first two stages. The long-standing BWA-GATK Best Practices, while widely adopted, present significant challenges in processing time, scalability for ultra-large cohorts, and storage requirements.

First, regarding individual variant calling, the standard BWA-GATK pipeline requires approximately 30 hours to process a 30X Whole Genome Sequencing (WGS) sample from FASTQ to VCF^1,2^. To address this, several accelerated suites have emerged. Heterogeneous computing solutions, such as MegaBOLT^2^ and DRAGEN^3^, leverage FPGA-based custom circuitry to consolidate computational instructions. Others, like NVIDIA Parabricks, utilize GPU acceleration. While these tools can reduce runtime to under 30 minutes, they necessitate specialized hardware, rendering existing CPU-only clusters obsolete and substantially increasing infrastructure costs.

Second, as cohort sizes reach the scale of 10^5 to 10^6 samples, conventional tools often encounter out-of-memory (OOM) errors on standard hardware. Existing solutions attempt to mitigate this through different architectures: GLnexus^4^ utilizes a sharding strategy with GVCF inputs but lacks the flexibility to perform joint calling on specific genomic intervals. Alternatively, GraphTyper^5^ adopts a pangenome strategy using BAM files as direct input, which bypasses GVCF generation but exacerbates storage pressures—especially since individual GVCF/VCF files remain essential deliverables for most researchers.

Finally, storage remains a critical bottleneck. A single 30X WGS sample generates 40–60 GB of GZIP compressed FASTQ data and 5–8 GB of GVCF data. For a cohort of 100,000 individuals, this translates to 4–6 PB of raw data and 500–800 TB of variant files, excluding backups.

To address these multifaceted challenges, we developed DCS Tools. Our design philosophy was centered on three pillars: acceleration without accuracy loss, hardware versatility, and storage optimization. By implementing hardware-level optimizations such as Single Instruction, Multiple Data (SIMD) and improving memory-resident data structures, DCS Tools eliminates the redundant input and output (I/O) operations inherent in traditional multi-step pipelines.

## System Implementation

### The DCS Tools Architecture

The DCS Tools suite is modularly structured into three functional pillars: a streamlined pipeline for individual variant calling (from FASTQ to VCF), an ultra-scalable joint calling engine, and a set of specialized genomic data compression tools(Table 1).

**Table 1.** Modules in DCS Tools.

| Module Name | Functionality |
| --- | --- |
| aligner | Quality control (QC), alignment, sorting, and duplicate marking |
| bqsr | Base quality score recalibration |
| variantCaller | Variant discovery (similar to GATK HaplotypeCaller) |
| genotyper | GVCF to VCF conversion |
| report | Generation of analysis reports |
| jointcaller | Joint variant calling with support for million-scale cohorts |
| SeqArc | Efficient and lossless FASTQ compression tool |
| VarArc | Efficient and lossless VCF compression tool |

### From FASTQ to Variant Discovery

DCS Tools implements a high-performance integrated pipeline tailored for large-scale population genomics. The architectural core focuses on instruction-level optimization of computationally intensive logic and the minimization of the entire I/O path, enabling exceptional execution efficiency on generic CPU architectures that significantly outperforms traditional toolchains.

#### 1. Integrated Alignment Engine (**aligner**)

In conventional bioinformatics pipelines, Quality Control (QC), sequence alignment, coordinate sorting, and duplicate marking are typically executed by discrete tools (e.g., fastp, BWA, Samtools, and Picard). This fragmented approach generates massive intermediate files, leading to severe disk I/O bottlenecks.

Consolidated Execution Stream: The aligner module integrates these disparate stages into a unified execution flow. Once sequence mapping is completed in-memory, the data is immediately channeled into the sorting and duplicate-marking logic, drastically reducing temporary disk footprint.

Flexible Memory Modes: For large-scale genomes or memory-constrained environments, DCS Tools offers a “Low Memory Index” mode. By optimizing the index storage structure, a 30X WGS analysis can be completed using approximately 50 GB of RAM in a 32-thread environment, ensuring superior compatibility with cloud instances and standard servers.

#### 2. High-Speed Base Quality Score Recalibration (**bqsr**)

The bqsr module corrects systematic biases inherent in the sequencing process. To optimize storage efficiency and minimize redundant I/O, DCS Tools adopts a streamlined approach by decoupling it from BAM file generation. Instead of producing a massive, recalibrated BAM file, the suite generates a compact recalibration information table. This table is then utilized directly as a input for subsequent variant calling modules. This design significantly reduces the cumulative output volume and mitigates the disk space overhead typically associated with large-scale genomic workflows.

#### 3. Robust Variant Detection and Genotyping (**variantCaller & genotyper**)

The variant discovery phase is handled by variantCaller, which identifies single nucleotide polymorphisms (SNPs) and small insertions/deletions (Indels) from the aligned sequences.

We implemented a variant discovery module in C++ that replicates the core logic of HaplotypeCaller, with a heavy emphasis on fine-grained parallelization to achieve significant computational acceleration.

Broad Taxonomic Applicability: Beyond standard human diploid analysis, variantCaller supports custom ploidy parameters, enabling efficient processing of complex polyploid genomes in plant and animal research.

Scalable GVCF Path for Large Cohorts: The module generates standard GVCF files by default, providing a seamless transition to the jointcaller module for subsequent million-scale cohort analysis.

#### 4. CPU-Centric Acceleration and Performance

The acceleration core of DCS Tools lies in “Engineering Refinement” rather than “Hardware Dependency.” By eschewing reliance on expensive GPU or FPGA hardware and focusing on sophisticated thread scheduling, cache optimization, and maximum utilization of modern CPU throughput, DCS Tools achieves ultra-high-speed processing. This approach substantially lowers the computational time and financial costs associated with population-scale genomic studies.

### Joint calling

DCS Tools integrates **DPGT**^6^ which parallelizes the joint calling process by partitioning on two dimensions: the sample dimension and the genome position dimension. The input of DPGT is a list of gVCF files, which can be generated by GATK[6,13], Sentieon[8] or Illumina DRAGEN[9], and a reference fasta file. First, DPGT combines gVCF headers of all samples. Next, the whole genome region is divided into many non-overlapping windows. In each window, DPGT performs variant sites finding, variants combining and genotyping. Finally, DPGT concatenates genotyped VCF files into one result VCF file.

### Data compression

The software implements two adaptive lossless compression modules for FASTQ(**SeqArc**, manuscript in preparation) and VCF(**VarArc**, manuscript in preparation). The FASTQ module structurally splits identifiers into fields, encoding numeric components with differential and dictionary coding; reads are preferentially aligned to reference using a sliding-window index, with unmatched segments entropy-coded by a high-order Markov model; quality scores are compressed via context-based prediction combined with arithmetic and run-length encoding. The VCF module reorders genotype matrices to reduce Hamming distances, employs columnar storage with pattern-based numeric, run-length, and dictionary coding, and both pipelines are finalized by a unified entropy-coding backend.

### Testing

We conducted a comprehensive evaluation of DCS Tools’ performance and accuracy using multiple gold-standard datasets. The benchmarking environment primarily consisted of Alibaba Cloud ecs.i4g.8xlarge instances (32-core CPU, 128 GB RAM, SSD storage) and local high-performance computing (HPC) clusters.

### Individual WGS Pipeline: Performance and Accuracy

We selected benchmark samples from the Genome in a Bottle (GIAB) consortium (HG001, HG002, HG003, and HG005), covering two major sequencing platforms: Illumina NovaSeq and MGI DNBSEQ, with sequencing depths ranging from 30X to 40X.

Execution Efficiency: Experimental results demonstrate that DCS Tools completes the end-to-end analysis from raw FASTQ to VCF in just 1.79 to 2.36 hours(Table 2). The alignment and duplicate marking stage (aligner), which is the most computationally intensive phase, accounts for approximately 65%–70% of the total runtime(Table 3). Both the Base Quality Score Recalibration (bqsr) and variant calling (variantCaller) modules exhibited exceptional parallel efficiency. Compared to the BWA-GATK Best Practices, DCS Tools achieves an approximate 10-fold acceleration under identical hardware configurations.

**Table 2.** Performance benchmarks of DCS Tools on WGS data (32 threads)

| Sample | Default configuration |  | Low Memory Index mode |  |
| --- | --- | --- | --- | --- |
|  | Runtime(h) | Max RSS(G) | Runtime(h) | Max RSS(G) |
| HG001 30X NovaSeq | 1.79 | 100 | 1.97 | 48 |
| HG002 30x NovaSeq | 2.15 | 112 | 2.31 | 58 |
| HG003 30x NovaSeq | 1.89 | 120 | 2.00 | 54 |
| HG005 40X DNBSEQ | 2.36 | 120 | 2.84 | 64 |

**Table 3.** Runtime of individual modules on HG001 30X NovaSeq (32 threads)

| Module | Runtime(h) |
| --- | --- |
| aligner | 1.21 |
| bqsr | 0.09 |
| variantCaller | 0.47 |
| genotyper | 0.02 |
| total | 1.79 |

Resource Optimization: In the “Low Memory Index” mode, the peak memory consumption of aligner is significantly reduced from over 100 GB to 48–64 GB, with only a marginal increase in runtime (approx. 10%–15%). This demonstrates the suite’s flexibility across diverse hardware environments with varying memory capacities.

Accuracy and Concordance: By benchmarking against the official GATK pipeline, DCS Tools achieved near-perfect concordance in SNP and Indel detection(Figure 1). The results meet the stringent accuracy standards required for both clinical diagnostics and advanced scientific research.

**Figure 1.**
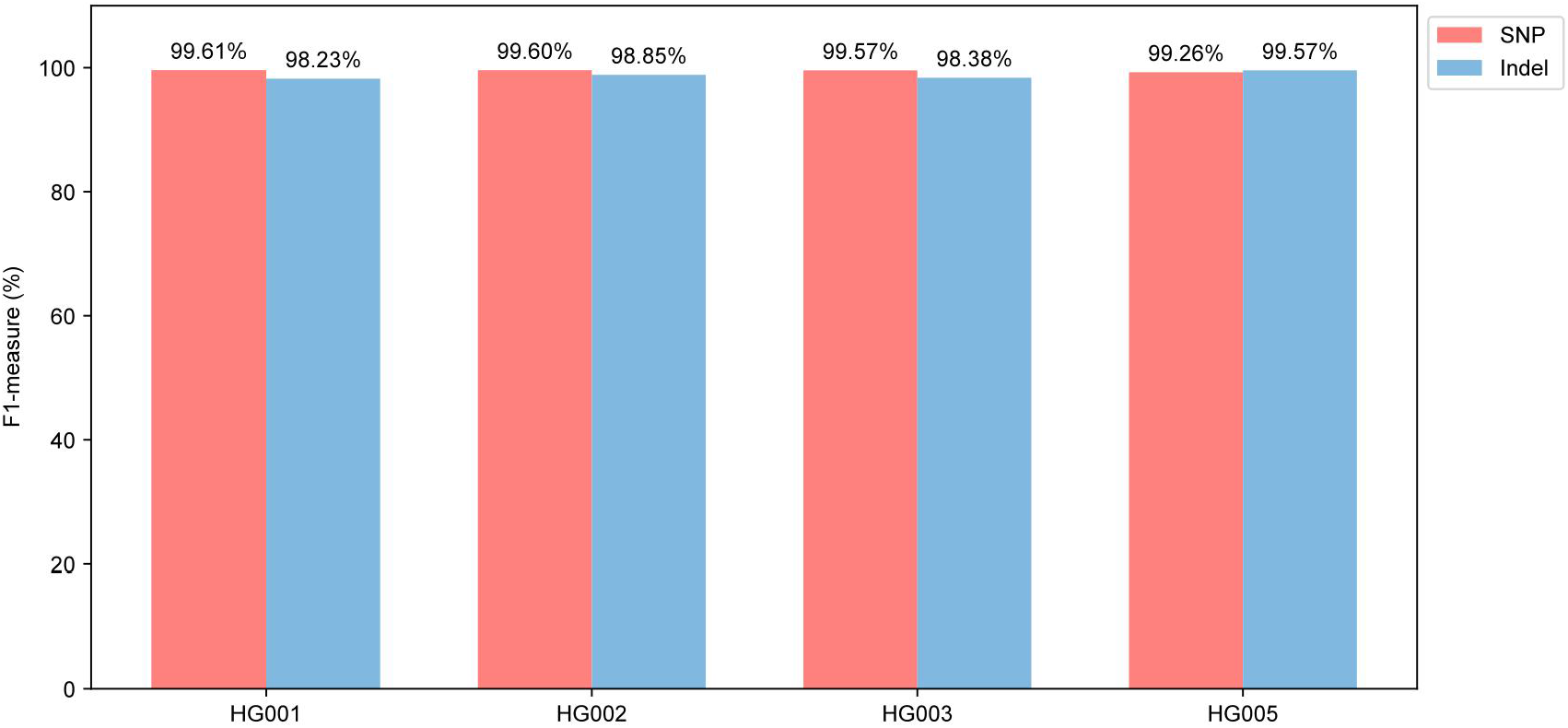
WGS Variant Calling Accuracy

### Large-scale Population Joint Variant Calling

To evaluate the scalability of the jointcaller module(DPGT), we conducted stress tests on cohorts ranging from ten thousand to hundreds of thousands of samples.

Processing Capacity: Benchmarking on a single 256-core server showed that jointcaller can process 10,000 samples in approximately 83 hours. In a real-world ultra-large-scale application, utilizing a distributed cluster of 300 nodes (each with 32 cores), DCS Tools successfully completed joint calling for 470,000 samples within 56 days(Table 4).

**Table 4.** Joint Calling Performance Across Different Cohort Sizes.

| Dataset | Size | Number of Processes(6 GB RAM per process) | Core-Hours | Runtime |
| --- | --- | --- | --- | --- |
| 1KGP | 2504 | 256 | 6963 | 27.2 h |
| Internal 10K | 9165 | 256 | 21376 | 83.5 h |
| Internal 52K | 52000 | 4400 | 407760 | ~6 d |
| Internal 105K | 105000 | 3675 | 1026155 | ~19 d |

Technical Advantages: This performance is attributed to the implementation of a proprietary linear index optimization (tbi2lix) and highly efficient operator parallelization. These innovations effectively mitigate the I/O amplification issues typically encountered during the stacking of massive GVCF files, supporting seamless scaling to million-sample cohorts.

### Data Compression Efficiency

To address the escalating costs of genomic data storage, we evaluated two specialized modules: SeqArc (for FASTQ) and VarArc (for VCF/GVCF).

FASTQ Compression (SeqArc): Compared to the industry-standard GZIP format, SeqArc’s “Single Reference” mode demonstrates a superior compression ratio, reducing file sizes to 1/4 or 1/5 of the original GZIP files(Table 5). Furthermore, its decompression speed allows for direct streaming into downstream analysis pipelines, optimizing the balance between storage costs and computational throughput.

**Table 5.** Compression Performance on FASTQ files with SeqArc.

| File(Size) | Runtime of compression | Compression Ratio (relative to GZIP input) | Runtime of decompression on .arc |
| --- | --- | --- | --- |
| HG001 r1(27G) | 27m 24s | 5.17 | 14m 32s |
| HG001 r2(30G) | 29m 14s | 4.10 | 14m 25s |

VCF/GVCF Compression (VarArc): VarArc excels in compressing variant output files. For individual GVCFs, it reduces file volume to approximately 1/3 of the GZIP size(Table 6). For population-scale VCFs (e.g., 100,000 samples), it achieves a size reduction to 1/2 of the GZIP equivalent(Table 7).

**Table 6.** Performance Comparison for Single-Sample GVCF with VarArc.

| GZIP size | comp_size(GB) | Compression Ratio | comp_Real Time(s) | decomp_Real Time(s) |
| --- | --- | --- | --- | --- |
| 51.6 | 15.52 | 3.3 | 4141.1 | 1213.4 |

**Table 7.** Compression Performance on Population-Scale VCF Files with VarArc.

| Data | File A | File B | File C |
| --- | --- | --- | --- |
| GZIP size(GB) | 15 | 25.9 | 60.7 |
| Samples | 6w | 10.5w | 10.5w |
| Variants | 8w | 8w | 524w |
| Fields Included | GT, AD, DP, etc. |  | GT only |
| Comp_size(GB) | 6.34 | 11.26 | 53.79 |
| Compression Ratio | 2.36 | 2.3 | 1.13 |
| Comp_RealTime(s) | 1143 | 1889 | 13772 |
| Decomp_RealTime(s) | 319 | 561 | 3864 |

Data Integrity: All compression modules have undergone rigorous bit-to-bit verification. Results confirm that the restored data is identical to the original input, ensuring the highest level of rigor for bioinformatics analysis.

## Future directions

The current version of DCS Tools primarily focuses on variant discovery based on linear reference genomes. To further enhance detection sensitivity and specificity—particularly in complex genomic regions—we are actively developing a pangenome-based variant calling engine. By incorporating graph-based genomic representations, this upcoming module aims to overcome the inherent biases of linear references and provide superior accuracy for diverse populations.

Additionally, to complete our high-performance data management ecosystem, we plan to release a specialized BAM compression tool in the near future. This addition will complement our existing SeqArc (FASTQ) and VarArc (VCF/GVCF) modules, offering researchers a comprehensive, end-to-end solution for minimizing the storage footprint across the entire genomic analysis pipeline.

## Availability and requirements

Software: https://tools.dcs.cloud/

Operating Systems: Linux (CentOS, Ubuntu, Debian, EulerOS).

Architectures: x86_64, ARM64 (aarch64).

## Data Availability

The Whole Genome Sequencing (WGS) benchmarks in this study were conducted using datasets from the Genome in a Bottle (GIAB) consortium, while data for other components were obtained from internal projects.

GIAB: ftp://ftp-trace.ncbi.nlm.nih.gov/giab/ftp/release/

## Authors’ contributions

Yong Zhang supervised the research. Yong Zhang, Shengkang Li, Chun Gong,Yuxin Chen, Xin Li and Qi Yang conceptualized the software. Dong Yuan, Chun Gong, Zijian Zhao, Qi Yang, and Ruiwen Wan implemented the software code. Qi Yang, Ruiwen Wan, Yuxin Chen and Zijian Zhao analyzed the data. Chun Gong contributed to the writing of the manuscript. All the authors read and approved the final manuscript.

## Acknowledgements

We would like to thank DCS Cloud (https://cloud.stomics.tech/) for providing the computational resources and software support necessary for this study.

## Notes

### Competing Interest Statement

The authors have declared no competing interest.

### Summary of Updates

add an author who contributed to the design of the software product

